# Colistin heteroresistance and the involvement of the PmrAB regulatory system in *Acinetobacter baumannii*

**DOI:** 10.1101/307132

**Authors:** Yannick Charretier, Seydina M. Diene, Damien Baud, Sonia Chatellier, Emmanuelle Santiago-Allexant, Alex van Belkum, Ghislaine Guigon, Jacques Schrenzel

**Affiliations:** Genomic Research Laboratory, Service of Infectious Diseases, Geneva University Hospitals, Geneva, Switzerland.; Innovation Unit, bioMérieux SA, La Balme Les Grottes, France.; Innovation Unit, bioMérieux SA, Marcy l’Etoile, France.; Data Analytics Unit, bioMérieux SA, La Balme Les Grottes, France.

**Author notes:** Address correspondence to Yannick Charretier,. Present address: Seydina M. Diene, Aix-Marseille Université, MEPHI, IHU-Méditerranée Infection, Marseille, France.

## Abstract

Multidrug-resistant *Acinetobacter baumannii* infection has recently emerged as a worldwide clinical problem and colistin is increasingly being used as last resort therapy. Despite its favorable bacterial killing, resistance and heteroresistance to colistin have been described. Mutations in the PmrAB regulatory pathway have been already associated with colistin resistance whereas the mechanisms for heteroresistance remain largely unknown. The purpose of the present study is to investigate the role of PmrAB in laboratory-selected mutants representative of global epidemic strains. During brief colistin exposure, colistin resistant and colistin heteroresistant mutants were selected in a one-step strategy. Population Analysis Profiling (PAP) was performed to confirm the suspected phenotype. Upon withdrawal of selective pressure, compensatory mutations were evaluated in another one-step strategy. A trans-complementation assay was designed to delineate the involvement of the PmrAB regulatory system using qPCR and PAP. Mutations in the PmrAB regulatory pathway were associated with colistin resistance and colistin heteroresistance as well. The transcomplementation assay provides a proof for the role played by changes in the PmrAB regulatory pathway. The level of colistin resistance is correlated to the level of expression of *pmrC*. The resistance phenotype was partially restored since the complemented strain became heteroresistant. This report shows the role of different mutations in the PmrAB regulatory pathway and warns on the development of colistin heteroresistance that could be present but not easily detected with routine testing.

## Introduction

*Acinetobacter baumannii* has emerged as an opportunistic and medically important pathogen causing a broad range of nosocomial infections, particularly amongst the most vulnerable patients in the intensive care units. The most frequent infections are ventilator-associated pneumonia and bacteremia, both of which leading to high morbidity and mortality (1). *A. baumannii* is intrinsically resistant to several classes of antimicrobial agents and readily acquires resistance to other classes leading to multidrug-resistant (MDR) *A. baumannii* isolates. Antimicrobial resistance provides them a selective ecological advantage that drives the clonal expansion of certain *A. baumannii* lineages such as the three international clonal complexes I, II and III (2).

Divalent cations bridging between lipopolysaccharide (LPS) molecules stabilize the membrane through balancing the electrostatic net charges (3). This unique composition of the outer membrane makes the cell susceptible to cationic antimicrobial peptides (CAMPs). Polymyxin B and polymyxin E (colistin), biosynthesized by some strains of *Paenibacillus polymyxa*, are two FDA-approved antibiotics of the CAMPs class that are increasingly used as the ultimate therapeutic agent in MDR Gram-negative bacteria such as *A. baumannii*. The mode of action of polymyxins is due to the amphipathic nature of these compounds. The positively charged cyclic group is thought to compete with divalent cations for binding to LPS while the acyl chain is thought to disrupt both the outer and the inner membranes (4, 5). To survive to these compounds, *A. baumannii* has developed two main antimicrobial resistance mechanisms: the loss of lipid A or its modification through modifying enzymes (6–8). The former mechanism, which overthrows a dogma that LPS is essential in Gram-negative bacteria, is due to the inactivation of downstream lipid A biosynthetic genes. This mechanism that is rare in clinical medicine confers a high level of colistin resistance. The loss of lipid A leads in return to a growth defect, higher fitness costs, virulence attenuation and hypersensitivity to other classes of antibiotics (6, 9, 10). The second mechanism is mediated by the operon *pmrCAB*. Genetic changes in the PmrAB regulatory system lead to the activation of *pmrA, pmrB* and *pmrC*, a phosphotransferase, which leads to the addition of phosphoethanolamine to lipid A (7, 8). Interestingly, *A. baumannii* may harbor a pmrC-like gene named *eptA* whose expression is increased in colistin resistant strains (11). This mechanism reduces net LPS negative charge leading to a low-to-medium level of colistin resistance. The impact on virulence and fitness depends on the underlying mutations and could range from low to apparently unaffected (9, 10).

Colistin heteroresistance has been described (12). Colistin heteroresistance has been associated with prior colistin exposure (13). This phenomenon is underappreciated due to the use of different diagnostic methods and standards. A standard definition of heteroresistance has been recently determined (14). Heteroresistant bacteria are defined as a single-species bacterial population displaying a heterogeneous response to an antibiotic, i.e. when the difference between the lowest concentration exhibiting maximum inhibition and the highest non-inhibitory concentration ranges above 8 fold. The reference method to define microbial response to an antibiotic is the Population Analysis Profiling (PAP). Clinically, the situation may be critical when the majority of the heterogeneous population is susceptible to an antibiotic and a subpopulation is resistant. This may lead to therapeutic failures due to susceptibility categorization errors. To date, no mechanism has been described for colistin heteroresistance in *A. baumannii*.

The purpose of the present study is to investigate the role of PmrAB in laboratory-selected mutants representative of clinical strains. During brief colistin exposure, colistin resistant and colistin heteroresistant mutants were selected in a one-step strategy. Upon withdrawal of selective pressure, compensation mechanisms were evaluated in a one-step strategy. PAP was performed to confirm the suspected phenotype. A trans-complementation assay was designed to delineate the involvement of PmrAB regulatory system using qPCR and PAP assessments.

## Results

### Mutations associated with colistin resistance

Following exposition to colistin for 24h, phenotypically resistant mutants were obtained in all clones. SNPs and indels could appear in the PmrAB regulatory pathway. Mutants derived from clinical isolates (Table 1) and obtained at extreme concentrations were further sequenced in the *pmrCAB* operon to reveal putative changes. Based on targeted sequencing, two mutants were found for Ab1, four mutants were found for Ab2 and only one mutant for Ab3 (Table 2). A unique mutant of strain Ab2 contained a mutation in the response regulator domain of the *pmrA* gene, while the three other were mutated in the *pmrB* gene. No changes were observed in the *pmrC* gene. Mutants of strain Ab3 displayed a unique mutation whatever the concentration used for the selection, whereas mutants of strain Ab2 harbored a mixed and more diverse population of mutants for each selection process. Mutants of strain Ab1 displayed an intermediate situation with a major mutant for each colistin concentration used for selection. Mutants in the histidine-kinase domain of the *pmrB* gene had the highest MICs and became colistin resistant. In contrast, other mutants were still susceptible to colistin despite the screening performed at 10 mg/L. Amino acid substitution in the PmrAB regulatory pathway is the prime suspect for colistin resistance but other genetic changes could be supposed. To address this hypothesis, two pairs (Ab1/Ab1-8-R1, Ab3/Ab3-2-R1) and one trio (Ab2/Ab2-1-R4/Ab2-1-R4-S1) of strains were sequenced. Genome-wide analysis of this subset confirmed the mutations observed using Sanger sequencing of the *pmrCAB* operon as well as the absence of non-synonymous mutations outside of the *pmrCAB* operon (data not shown). Only a synonymous mutation was observed in a putative siderophore for Ab1-8-R1. Two different mutations were observed in a non-coding region for Ab2-1-R4-S1 and for Ab3-2-R1 respectively (data not shown).

**Table 1:** Clinical isolates used in this study.

| Isolates | Clinical source | ST<sup>UO</sup> | ST<sup>IP</sup> | CC<sup>UO</sup> / CC<sup>IP</sup> |
| --- | --- | --- | --- | --- |
| Ab1 | Blood culture | 449 | 20 | 109 / 1 |
| Ab2 | Rectal swab | 208 | 2 | 92 / 2 |
| Ab3 | Skin swab | 106 | 3 | 187 / 3 |

**Table 2:**
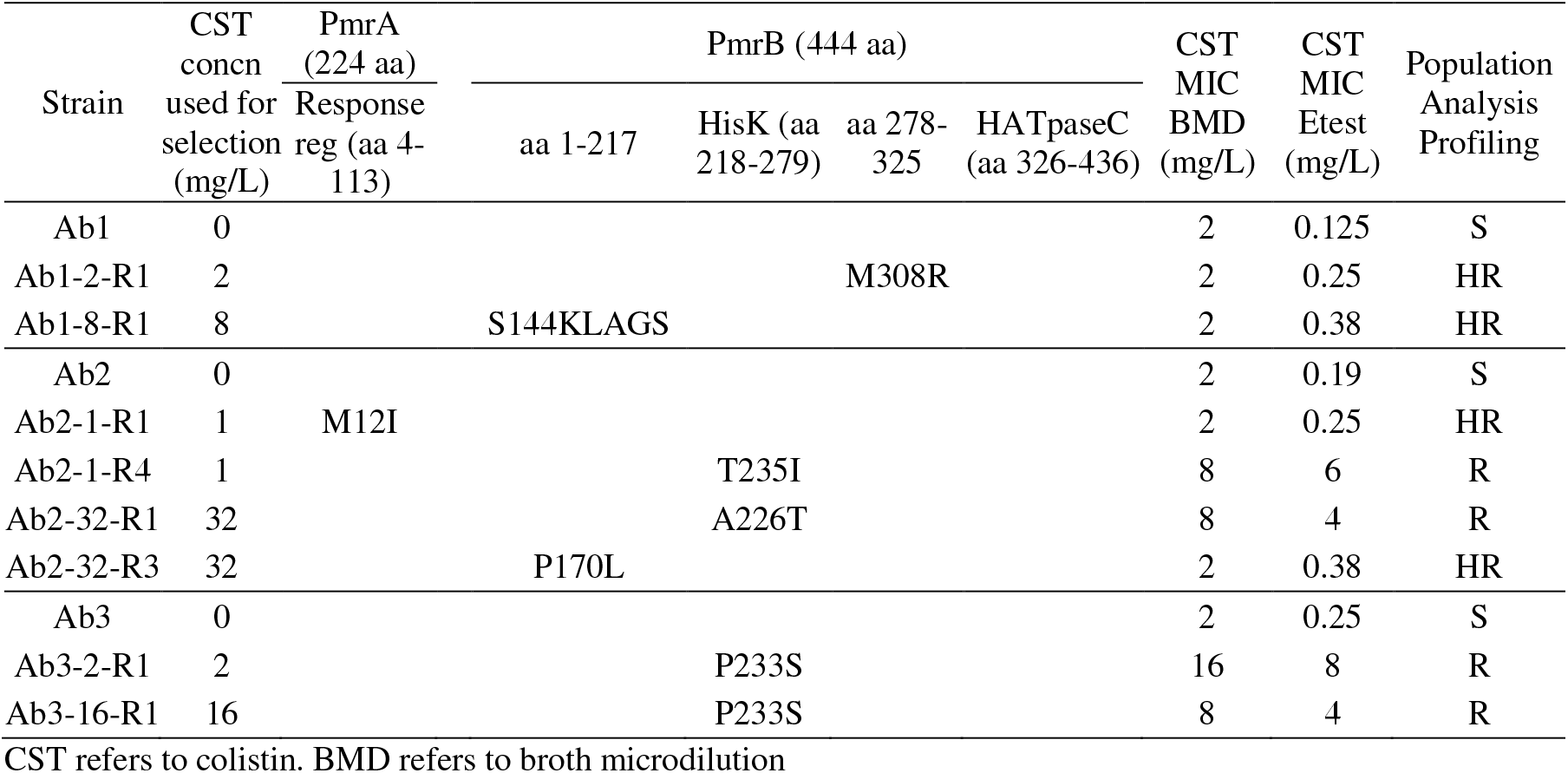
Mutational analysis of the PmrAB regulatory pathway

### Mutations associated with the loss of colistin resistance

The second aim was to assess whether compensatory mutations would occur using *in vitro* laboratory conditions and a one-step selection procedure. A single mutant, the one with the highest MIC, was screened for each clone. The loss of colistin resistance was observed through a compensatory mutation in Ab2-1-R4. An IS*Aba1* insertion sequence (1188-bp in length) was inserted after the proline 170 within the *pmrB* gene and generated a 9-bp duplication at the extremities of the insertion site. Despite many attempts to isolate Ab3-2-R1 (48, 54, 161, 297 and 576 individual colonies tested per screen), the screened sampling failed to highlight the loss of colistin resistance. The screening procedure was unsuitable for isolates Ab1-2-R1 and Ab1-8-R1 that were heteroresistant mutants.

### Mutants confirmed as resistant and heteroresistant to colistin

To confirm the character of colistin resistance of the mutants, PAPs were carried out. First, the response to colistin of the three clinical strains showed a homogeneous response to colistin in the susceptible range (Fig. 1A-C). Second, for many of the mutants, heteroresistance was suspected since MICs were similar to those of the parental strain. Mutants with an MIC of 2 or below exhibited heterogeneous responses (Fig. 1A-B) to colistin while mutants with an MIC of 8 or above exhibited a clearly resistant population (Fig. 1B-C). Mutants harboring mutations outside of the histidine-kinase domain of the *pmrB* gene were confirmed as heteroresistant mutants. As additional changes could be suspected for the resistant subpopulation, PAPs were performed on the resistant subpopulations for isolate Ab1-8-R1 and isolate Ab2-1-R1 that grew in agar plates containing 16 mg/L of colistin (data not shown). Both isolates remained heteroresistant although the proportion of resistance for isolate Ab2-1-R1 was larger (10^−2^) than the original one (10^−6^). With regard to the loss of colistin resistance, isolate Ab2-1-R4-S1 displayed a phenotypic reversion. It had the same profile as the susceptible parental strain (Fig. 1B). The insertion sequence IS*Aba7* may have inactivated the *pmrB* gene. If the sensor protein is inactivated, the regulator protein cannot be up-regulated, as well as the enzyme responsible for lipid A modification, the product of the *pmrC* gene.

**Figure 1:**
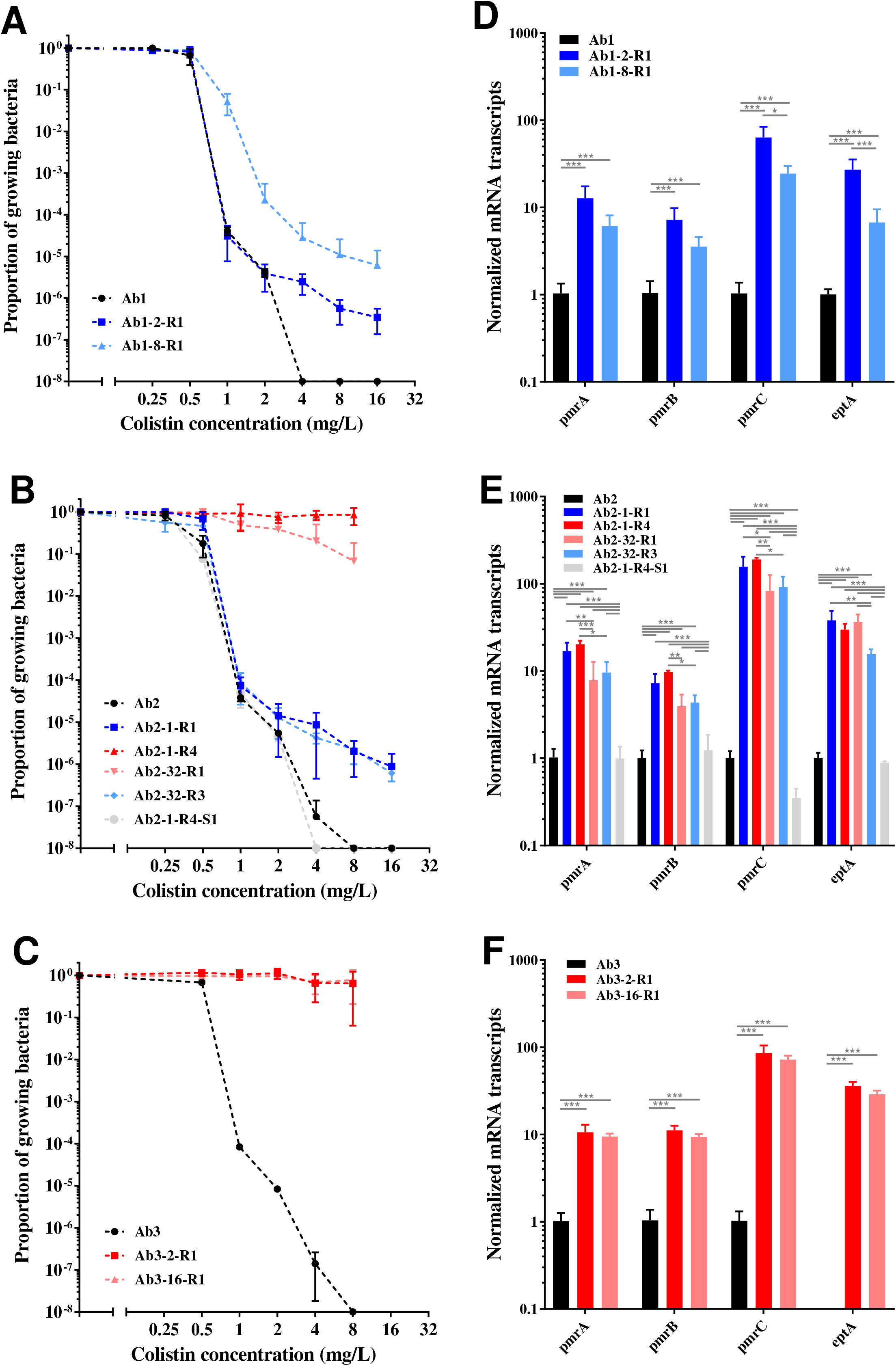
Response to colistin of resistant and heteroresistant mutants (A-C) and transcript levels of putative PmrA-regulated genes in sequentially derived isolates (D-F). Population analysis profiling (PAP) of isolates derived from Ab1 through Ab3 (A-C). PAP obtained by the spotting method (n=3). Dark grey curves and sticks refer to parental isolate, blue curves and sticks refer to heteroresistant mutants, red curves and sticks refer to resistant mutants and light grey curves and sticks refer to the mutant loosing colistin resistance. For the transcriptional experiment (D-F), data represent the mean from 3 independent experiments; error bars represent 1 SD. One-way analysis of variance with the Tukey multiple-comparison test was used to determine whether transcript levels from a gene differed among the strains. * refers to P < 0.05, ** refers to P < 0.01, *** refers to P < 0.001.

### Mutants associated with colistin resistance and heteroresistance

To verify whether there was an association between mutants and the PmrAB regulatory pathway, it was decided to quantify the level of transcripts of previously described genes in longitudinally derived isolates. The aim was to study if one of these genes had a distinct expression according to the susceptibility to colistin (susceptible, heteroresistant, resistant). As expected, the expression of the transcripts of the *pmrCAB* operon increased both, in colistin resistant and colistin heteroresistant mutants (Fig. 1D-F). Unexpectedly, the level of transcripts of *pmrC* were highly similar in colistin-resistant and colistin-heteroresistant strains derived from Ab2 (Fig. 1E). The role of colistin as an inducer of the expression of the transcripts of the *pmrCAB* operon was checked. Two stress concentrations of colistin (0.25 mg/L and 2 mg/L) were tested for Ab2 mutants and no induction was observed (data not shown). In isolate Ab2-1-R4-S1, the transcript level of the *pmrCAB* operon came back to the level observed in the parental clinical isolate confirming the role played by the insertion sequence within the *pmrB* gene. Furthermore, the transcript level of the *pmrC* gene remained even lower than the transcript level observed in the parental strain. The expression of the transcripts of *eptA* showed the same trend as that of the genes present in the *pmrCAB* operon except that the transcripts of *eptA* were not constitutively expressed in Ab3 (Fig. 1F).

### PmrAB mutants contributed to colistin resistance and colistin heteroresistance

To check if amino acid substitutions identified by Sanger sequencing contributed to colistin resistance or heteroresistance, isolate Ab2 was complemented *in trans* with the wild-type and mutated *pmrAB* genes. Complementation of isolate Ab2 with the wild-type *pmrAB* genes did not alter the susceptibility to colistin (Fig. 2A). Complementation with a heteroresistant mutant or a disrupted mutant did not alter the susceptibility to colistin (Fig. 2A). Colistin resistance was only partially restored when isolate Ab2 was complemented with *pmrAB_(Ab2-ı-R4)_*. Complementation with a resistant mutant resulted in colistin heteroresistance (Fig. 2A). The level of transcripts of the *pmrA* and *pmrB* genes in transformants complemented *in trans* with the wild-type and mutated *pmrAB* genes were equally upregulated (Fig. 2B). These results indicate that wild-type and mutated *pmrAB* genes were actually cloned with their own promoter. The level of transcripts of the *pmrC* gene was up-regulated for the resistant- and the heteroresistant-mutants complementation only. The highest levels were observed for the resistant-mutant complementation. The level of colistin resistance was correlated to the level of expression of *pmrC*. This result was consistent with a direct regulation with PmrA when amino acid substitution occurred in the PmrAB regulatory pathway. The same trend was observed for *eptA* except that the up-regulation was less marked and held for the resistant-mutant exclusively. Interestingly, the level of transcripts of the *pmrC* gene for the disrupted *pmrB* gene complementation was not lower than the level of transcripts observed in the parental strain (Fig. 2B).

**Figure 2:**
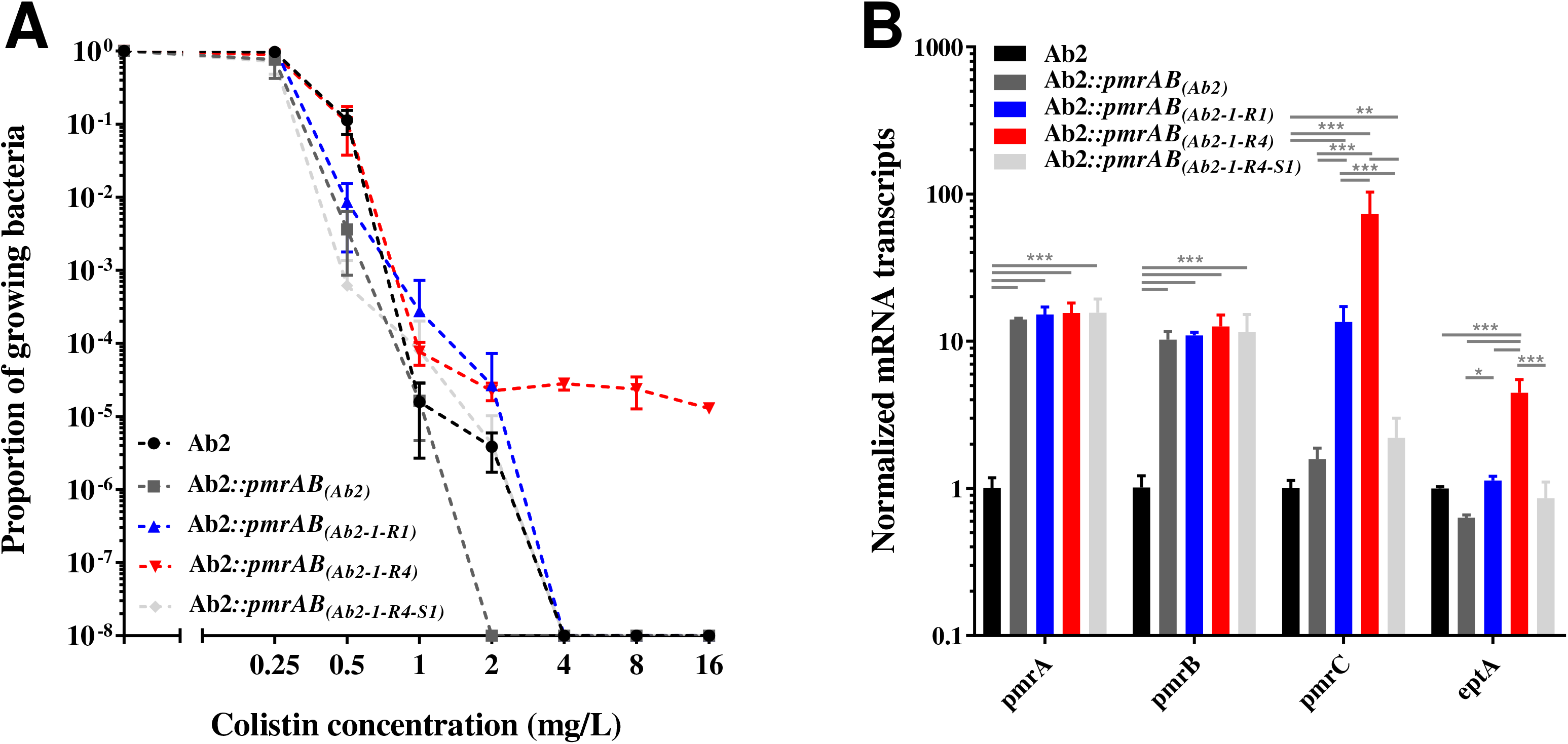
Response to colistin of trans-complemented transformants (A) and transcript levels of putative PmrA-regulated genes in trans-complemented transformants (B). Population analysis profiling (PAP) of transformants derived from trans-complemented Ab2 mutants (A). PAP obtained by the spotting method (n=3). Black curve and stick refer to parental isolate, dark grey curve and stick refer to a transformant complemented by the parental *pmrAB* genes from isolate Ab2. Blue curve and stick refer to a transformant complemented by the heteroresistant *pmrAB* genes from isolate Ab2-1-R1. Red curve and stick refer to a transformant complemented by the resistant *pmrAB* genes from isolate Ab2-1-R4. Light grey curve and stick refer to a transformant complemented by the *pmrAB* genes from isolate Ab2-1-R4-S1. For the transcriptional experiment (B), data represent the mean from 3 independent experiments; error bars represent 1 SD. One-way analysis of variance with the Tukey multiple-comparison test was used to determine whether transcript levels from a gene differed among the transformants. * refers to P < 0.05, ** refers to P < 0.01, *** refers to P < 0.001.

## Discussion

Heteroresistance is a phenomenon where subpopulations of seemingly isogenic bacteria exhibit a range of susceptibilities to a particular antibiotic. Heteroresistance can be intrinsic or acquired. In this study, colistin heteroresistance as well as colistin resistance were acquired through mutations in the PmrAB regulatory pathway. The trans-complementation assay demonstrated the role played by this two-component system and showed a direct correlation between expression of pmrC and colistin (hetero)resistance.

The one-step selection strategy generated both resistant and heteroresistant isolates. All mutations described in this study have been previously observed in clinical or laboratory strains except the two heteroresistant mutants derived from isolate Ab1 where a 12 base pair insertion or a M308R amino acid substitution were observed in *pmrB*. The fact that these two mutants had not been observed yet could be due to the process of selection for spontaneous mutation production chosen by the other authors (7, 8, 15, 16). The authors favored serial gradient strategies to promote stable mutants. As an example, Arroyo *et al*. were the first to identify the M12I mutation in the *pmrA* gene exhibiting a polymyxin B MIC of 4 mg/L, compared to the parental MIC of 0.5 mg/L. Their mutant was correctly classified as resistant and the authors did not perform PAP as a result (8). Their process for selection may increase the proportion of resistant subpopulations for this specific mutant as observed with our isolate making a heteroresistant mutant fully resistant to colistin. This characteristic may explain why they did not succeed to obtain knock-out for this mutant. Regarding the heteroresistant mutant derived from isolate Ab2 and presenting the P170L mutation, Pournaras *et al*. have related this mutation to colistin resistance before (MIC=32 mg/L) (17). Interestingly, Arroyo *et al*. have identified two clinical strains with mutations at that same position in the *pmrB* gene (8). The former clinical strain exhibited a different mutation, P170Q, leading to colistin resistance (MIC=16 mg/L). The latter clinical strain exhibited the same mutation but also the A80V mutation (MIC=128 mg/L). The detection of mutations appears currently not sufficient for inferring phenotypes. To support this observation, Park *et al*. have studied colistin resistance without mutations in known colistin resistance genes (16). Although the P233S mutation observed in Ab3 mutant seemed to be exclusively related to colistin resistance as demonstrated before (17–20), the discovery of a clinical colistin-revertant strain with compensatory mutations further restricts phenotype inference based on the genotype (18).

The level of colistin resistance are still low (1-4%) in *Acinetobacter* spp. (21, 22). The hypothesis of reduced virulence of resistant *Acinetobacter baumannii* strains was previously raised in the literature, among other possibilities (23). The possibility of compensatory mutations restoring the virulence in colistin-resistant bacteria was noticed (24) and actually observed in a case-study (25). To explain the low prevalence of colistin resistance, the hypothesis of heteroresistance could not be ruled out. Two animal-based studies proved that *K. pneumoniae* and *E. cloacae* displaying undetected colistin heteroresistance could lead to treatment failure in murine infection (26, 27). Heteroresistance might be an alternative to the costly colistin resistance. A transcriptome remodelling has been portrayed during infection and treatment and a convergent change was observed for clinical isolates with *pmrAB* mutations (28). Most isolates originated from patients treated with colistin. All of the isolates were susceptible to colistin at an MIC of 0.5 mg/mL and had a differential expression of *pmrC* that is consistent with heteroresistance. In addition, this study allowed to more strictly define the Pmr regulon in agreement with a precedent transcriptomic study (29). The PmrAB two-component regulatory system facilitates adaptation to diverse environments in many Gram-negative bacteria (30). The use of CAMPs may confer a cross-resistance to host CAMPs and the *A. baumannii* PmrAB two-component regulatory system could be the culprit (31). Heteroresistance due to *pmrAB* mutations may also have implications on immunogenicity. In *Francisella tularensis, flmK* leads to addition of galactosamine to lipid A and its inactivation provided protective immunity (32). The galactosamine modification of lipid A has been described in *Acinetobacter baumannii* as well (33).

Therapeutic concentrations of colistin range from 2 to 4 mg/L in patients (34). Concentrations to prevent the selection of resistant mutants are considerably higher than therapeutic concentrations (35). It is not surprising that the rate of heteroresistance can vary from 18.7% to 100% (36). Disk diffusion or MIC gradient strips assays are typically used as screening methods (14). They might miss the heteroresistant phenotype because of the low bacterial load used in these methods, a load that is inappropriate to detect the (low) frequency resistant subpopulation. In addition, colistin activity might be overestimated if tested in cation-adjusted Mueller-Hinton broths (as recommended) as compared to interstitial space fluid-like media possibly causing a shift from susceptible to resistant in *A. baumannii* (37). At the opposite, adsorption of polymyxins to plastics might overestimate colistin MICs which can be prevent by the use of low-protein-binding polypropylene (38). Although a study proved that reproducibility and robustness were superior using agar dilution compared to broth dilution methods (39). Both CLSI and EUCAST recommend the use of broth microdilution as the only valid method for MIC testing. A standardized protocol globally accepted for colistin susceptibility testing would be more than welcome. The clinical relevance of heteroresistance was not proven at that time impaired by the difficulties to measure the right MIC (40) and to detect heteroresistance *a fortiori* (14).

## Material and Methods

### Bacterial strains, growth conditions and susceptibility testing

A collection of 27 MDR *A. baumannii* isolates was collected at Geneva University Hospital (41). Multilocus Sequence Typing was performed to identify and select strains of different genetic backgrounds (42). Three clinical isolates (Ab1, Ab2, Ab3), belonging to the international clones I, II or III were randomly selected (Table 1). All strains were stored in glycerol stocks at −80°C. Subcultures were performed on Mueller-Hinton agar medium (Becton Dickinson AG, Allschwil, Switzerland) at 37°C for 18h in aerobic conditions unless described otherwise. Colistin susceptibility testing was done by broth microdilution using cation-adjusted Mueller Hinton broth medium (Becton Dickinson AG) according to the Clinical and Laboratory Standards Institute (CLSI). The CLSI and EUCAST breakpoints are the same for *A. baumannii* /colistin but the need to continue with the revision of the methodology for testing and breakpoints is still reaffirmed (40). To date, an isolate is susceptible to colistin when the MIC is ≤2 mg/L and resistant when above. MICs were also determined using Etest strips (bioMérieux SA, Marcy l’Etoile, France) on Mueller-Hinton agar plates (Becton Dickinson AG) inoculated with a 0.5 McFarland suspension.

### Selection of colistin-resistant and colistin-reversion mutants

Isolates Ab1, Ab2 and Ab3 were sub-cultured in 5 mL Lysogeny Broth (LB) (Becton Dickinson AG) containing colistin (Sigma-Aldrich, Saint Louis, United States) concentrations ranging between 0 to 32 mg/L for 24h at 37°C with shaking in 50 mL Falcon tubes (Corning Science Mexico, Tamaulipas, Mexico). In a one-step liquid-based culture, resistant mutants were isolated from LB plates containing 10 mg/L of colistin selected at extreme concentrations (n=4 per condition). Resistant mutants were stored as glycerol stocks at −80°C. Overnight cultures of resistant mutants were performed in 5mL LB medium at 37°C for 18h and a substantial number of clones (e.g. > 100) were isolated. The loss of colistin resistance was screened by comparing growth on LB in presence/absence of colistin. Mutants were isolated and saved (n=4 per condition).

### Bacterial lysis and Sanger sequencing of the *pmrCAB* operon

Single colonies picked from LB agar plates were suspended in 100μL Tris-EDTA buffer (Tris 10 mM and EDTA 1mM, pH8) containing acid washed glass beads (Sigma-Aldrich, Saint-Louis, MO). The cells were disrupted using an orbital shaker (Vortex Genius 3, IKA, Staufen, Germany) for 1 min and the lysate was centrifuged at 3500g for 3 min. Five μL of hundredfold diluted supernatant were directly used for PCR of the entire *pmrCAB* operon using previously described primers (Table 3). The PCR products were purified using QIAquick PCR purification kit (Qiagen, Hilden, Germany) following supplier instructions and eluted in 30 μL of pure water. Both strands were completely sequenced with Big Dye V3.1 on a 3730XL DNA Analyzer (Life Technologies, Carlsbad, CA) using the same primers as were used for amplification (Table 3).

**Table 3:**
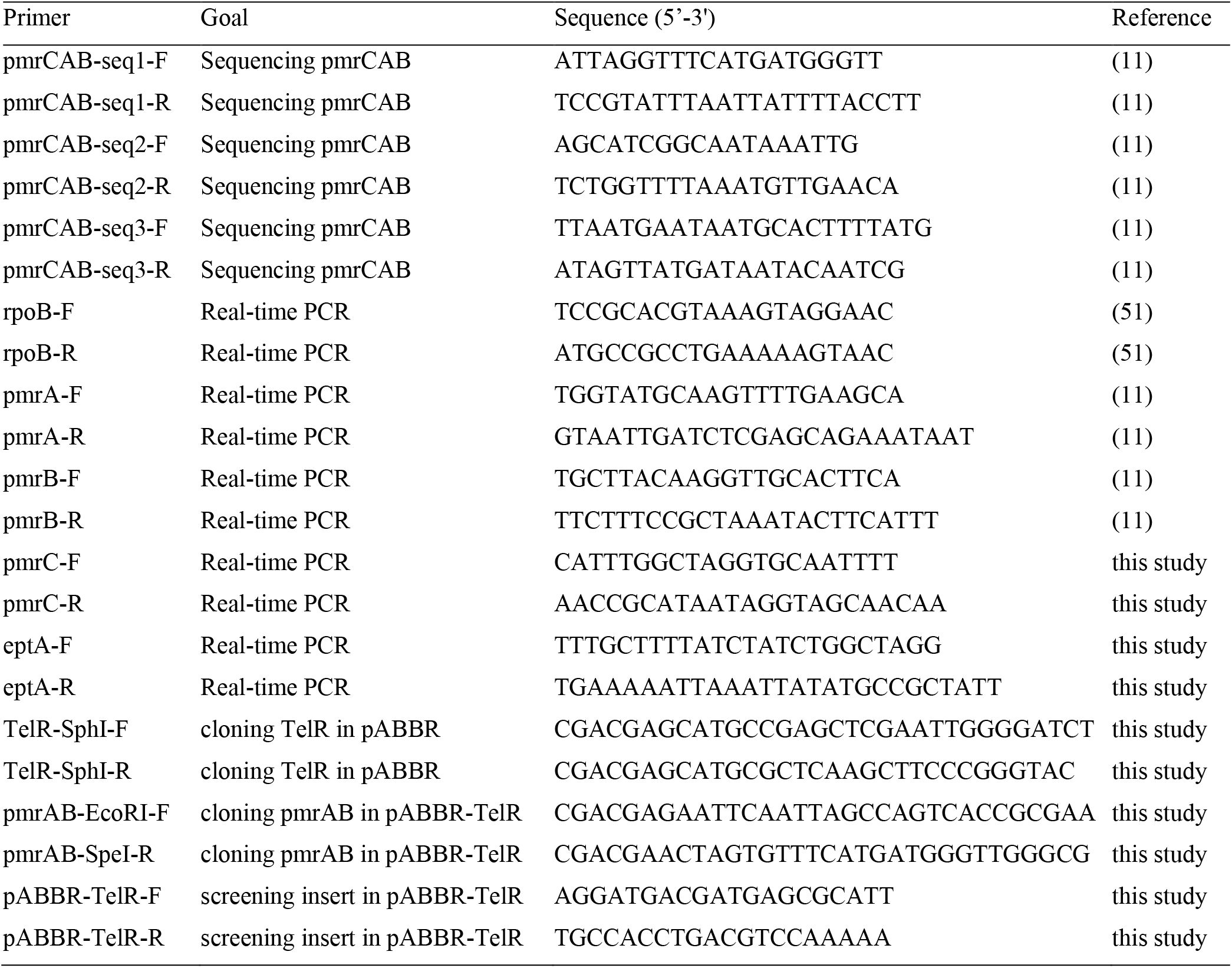
Primers used in this study

### Population Analysis Profiling (PAP)

Response to colistin in selected mutants was determined by PAP using a modified spotting method (43). Briefly, a suspension of each strain was adjusted to 2.0 McFarland and serially 10-fold diluted until 10^−7^. Both isolates Ab1 and Ab2, and their mutants were streaked on MH alone and on a two-fold series of colistin-containing plates from 0.25 to 16 mg/L. Isolate Ab3 and mutants were streaked on MH alone and on a two-fold series of colistin-containing plates from 0,5 to 8 mg/L. For each condition, four 10-μL droplets of the four highest concentrations were spotted on one plate and four 10-μL droplets of the lowest concentrations were spotted on a second plate. Colonies were counted after 24h incubation at 37°C. Three biological replicates were performed. Heteroresistance was considered when the antibiotic exhibiting the highest inhibitory effect was >8-fold higher than the highest non-inhibitory concentration (14).

### RT-qPCR experiments

The selected clinical isolates and mutants were grown to the mid-logarithmic phase in cation-adjusted Mueller Hinton broth. Suspensions were adjusted to an optical density of 0.01 at 595 nm and grown during five hours with shaking at 37°C before harvesting. Isolation of total RNA and synthesis of cDNA were performed as recently published (44) except that Superscript II^TM^ reverse transcriptase was used (Invitrogen, Carlsbad, CA). The primers for the PCR amplification of cDNA were retrieved from literature or designed using the primer3 program (45) (Table 3) and validated using the MIQE guidelines (46). An Mx3005P qPCR system (Agilent, Santa Clara, United States) was used for the quantification of cDNA. Triplicate PCRs were performed using the Brilliant III Ultrafast SYBR Green Q-PCR Master mix (Agilent). The thermal profile was performed according to supplier instructions. To correct for differences in the amount of starting material, the *rpoB* gene was selected as a reference gene. Results were presented as ratios of gene expression between the target gene and the reference gene, which were obtained according to Pfaffl *et al*. (47). Differences in the PCR efficiencies were corrected according to LinRegPCR program (48) and assessment of PCR efficiencies was based on the median of individual values calculated for each amplicon. Biological triplicates were performed. One-way analysis of variance with the Tukey multiple-comparison test was used to determine whether transcript levels from a gene differed among the isolates (P≥0.05).

### Genome sequencing

Whole genome sequencing of the 7 *A. baumannii* isolates was performed using Illumina high-throughput sequencing technology; the Ab1 pair (Ab1/Ab1-8-R1); the Ab2 trio (Ab2/Ab2-1-R4/Ab2-1-R4-S1); the Ab3 pair (Ab3/Ab3-2-R1). Extraction and purification of the genomic DNAs of the 7 isolates were achieved using an automated system (Nuclisens^®^ easyMag^®^ bioMerieux SA, Marcy l’Etoile, France). DNA quantity and purity in extracts were controlled using a fluorometer (Qubit Molecular Probes, Eugene, OR) and a spectrophotometer (ND-2000 NanoDrop Technologies, Wilmington, DE). Library preparation was performed with the Nextera XT DNA sample preparation kit (Illumina, San Diego, CA) and Nextera XT index kit (Illumina) following manufacturer’s recommendations. Library validation was performed on a 2100 Bioanalyzer (Agilent Technologies, Santa Clara, CA) to control the distribution of fragmented DNA. Sequencing runs of paired-end 2×250 cycles with the MiSeq Reagent kit v2 (500 cycles) (Illumina) on a MiSeq instrument (Illumina) was performed.

### Genome assembly and mutation detection

The genome was assembled using a version of the A5 pipeline called A5-miseq (49). Briefly, A5-miseq consists of three main stages: i) read quality filtering and error correction, ii) contig assembly and draft scaffolding with insert size, depth of coverage evaluation and error correction by read pairs mapping on contigs, iii) mis-assembly detection, conservative scaffolding and error correction by read pairs stringent mapping on contigs to generate the final consensus. The parental isolates were assembled in 50 to 77 scaffolds resulting in reference draft genomes, which were used to next mutation detection of resistant and compensatory mutants. Punctual mutations supported by at least three reads after read pairs mapping were kept for following interpretation.

### Trans-complementation assay

The TelR cassette from the pMo130-TelR plasmid was PCR amplified and cloned into the SphI site of pABBR_MCS to create the pABBR-TelR plasmid. The *pmrAB* genes from Ab2, Ab2-1-R1, Ab2-1-R4 and Ab2-1-R4-S1 were PCR amplified and cloned into the MCS of pABBR-TelR. Cloning was carried out in *E. coli* DH5α thermos-competent cells. The transformation protocol based on heat shock method was performed as recommended by the supplier (New England Biolabs, Ipswich, MA). The antibiotics kanamycin 7.5mg/L and ampicillin 100mg/L were added as needed for the selection. The complementation plasmids were transformed by electroporation (25uF, 1.8kV and 100 ohm in 0.2 mm cuvettes) as previously described (50). Transformants were selected in LB-agar plates supplemented with tellurite at 60 mg/L.

### Accession numbers

The genome sequences were deposited in the GenBank/ENA/DDBJ databases under accession nos. LNUX00000000 (Ab3), LNUY00000000 (Ab1), LNUZ00000000 (Ab2), LNVA00000000 (Ab3-2-R1), LNVB00000000 (Ab1-8-R1), LNVD00000000 (Ab2-1-R4-S1), LNVE00000000 (Ab2-1-R4).

